# Synthbar: A Lightweight Tool for Adding Synthetic Barcodes to Sequencing Reads

**DOI:** 10.1101/2025.05.30.657070

**Authors:** Jacob Morrison, Benjamin K. Johnson, Hui Shen

## Abstract

Preparation of single-cell sequencing libraries includes adding nucleotide barcodes to assist with pooling samples or cells together for sequencing. The popularity of droplet-based single-cell protocols has spurred the development of computational tools that expect the read structure of the assay to include a cell barcode (CB). Microwell plate-based protocols, such as the Switching Mechanism At the 5’ end of the RNA Transcript (SMART) single-cell RNA sequencing (scRNA-seq) family of methods, typically do not add a CB as part of the library preparation method as there is typically one cell per well and standard unique dual indices are sufficient for multiplexing. While several tools exist to manipulate and parse varying single-cell read structures, no tool is currently available to easily add synthetic CBs to enable use of computational tooling that expects the presence of a CB, such as STARsolo, zUMIs, and Alevin. Synthbar fills this gap as a lightweight tool that is assay agnostic, can add user-defined CBs, and modify read structures.

**Availability and Implementation:** Source code and binaries are freely available at https://github.com/jamorrison/synthbar under the MIT license. Synthbar is implemented in C and is supported on macOS and Linux.

**Supplementary information:** Supplementary data are available at the end of the document.

## Introduction

It is common practice in single-cell whole genome (DNA) and RNA (cDNA) sequencing to attach additional nucleotide sequences in order to separate biologically relevant reads from a pooled/multiplexed library or account for library preparation biases. These sequences typically are either a cell barcode (CB), which distinguishes one cell from another, or a unique molecular index (UMI), which identifies distinctive original DNA/cDNA fragments. Droplet-based single-cell RNA-seq protocols often include both a cell barcode and a UMI (e.g., 10x Genomics), whereas plate-based protocols may only utilize one or the other. For example, SMART-seq-total (4), FLASH-seq (3), MATQ-seq (12), Smart-seq3xpress (2), SMART-seq2 (10), and STORM-seq (5), among other plate-based single-cell RNA-seq (scRNA-seq) protocols, and even some bulk protocols, include only a UMI, but no cell barcode. With the popularity of droplet-based methods and data, many tools for analyzing scRNA-seq with UMIs either require a CB (e.g., STARsolo (6), zUMIs (9), and Alevin (14)) or follow a multi-step process of alignment, tagging, and de-duplication to analyze the data (e.g., UMI-tools (13)).

Tool kits such as splitcode (15), seqtk (https://github.com/lh3/seqtk), and seqkit (11) are powerful computational tools to manipulate FASTA/FASTQ files for a variety of read-parsing operations. However, to the best of our knowledge, no tool currently allows for the addition of user-defined synthetic cell barcodes to read structures, and manipulate the new structure to meet expected inputs for tools requiring a cell barcode (or UMI). While it is possible to accomplish this task using combinations of existing tools, the required domain knowledge limits accessibility to perform this operation. Moreover, the use of configuration files that a user must annotate for a toolkit to work can also limit general accessibility.

To overcome this hurdle, we introduce synthbar, a commandline tool for adding synthetic cell barcodes, UMIs, or arbitrary nucleotide sequences to FASTQ reads. Synthbar handles reads from both uncompressed or gzip compressed FASTQ files and runs in O(*n*) time.

## Description

Synthbar is written in C and utilizes klib (https://github.com/attractivechaos/klib). Klib’s kseq provides native reading of compressed and uncompressed FASTQ files, as well as from standard input. Klib’s kstring implementation provides a dynamic C string type that can be altered on the fly, which allows synthbar to manipulate read structures in a lightweight, flexible manner. Further, synthbar provides several command line options, such as providing a user-defined nucleotide sequence of any length and composition to serve as the cell barcode or synthetic UMI, modifying the read structure to accommodate the inserted sequence, or removing other technical sequences (Figure 1A–D).

**Fig. 1.**
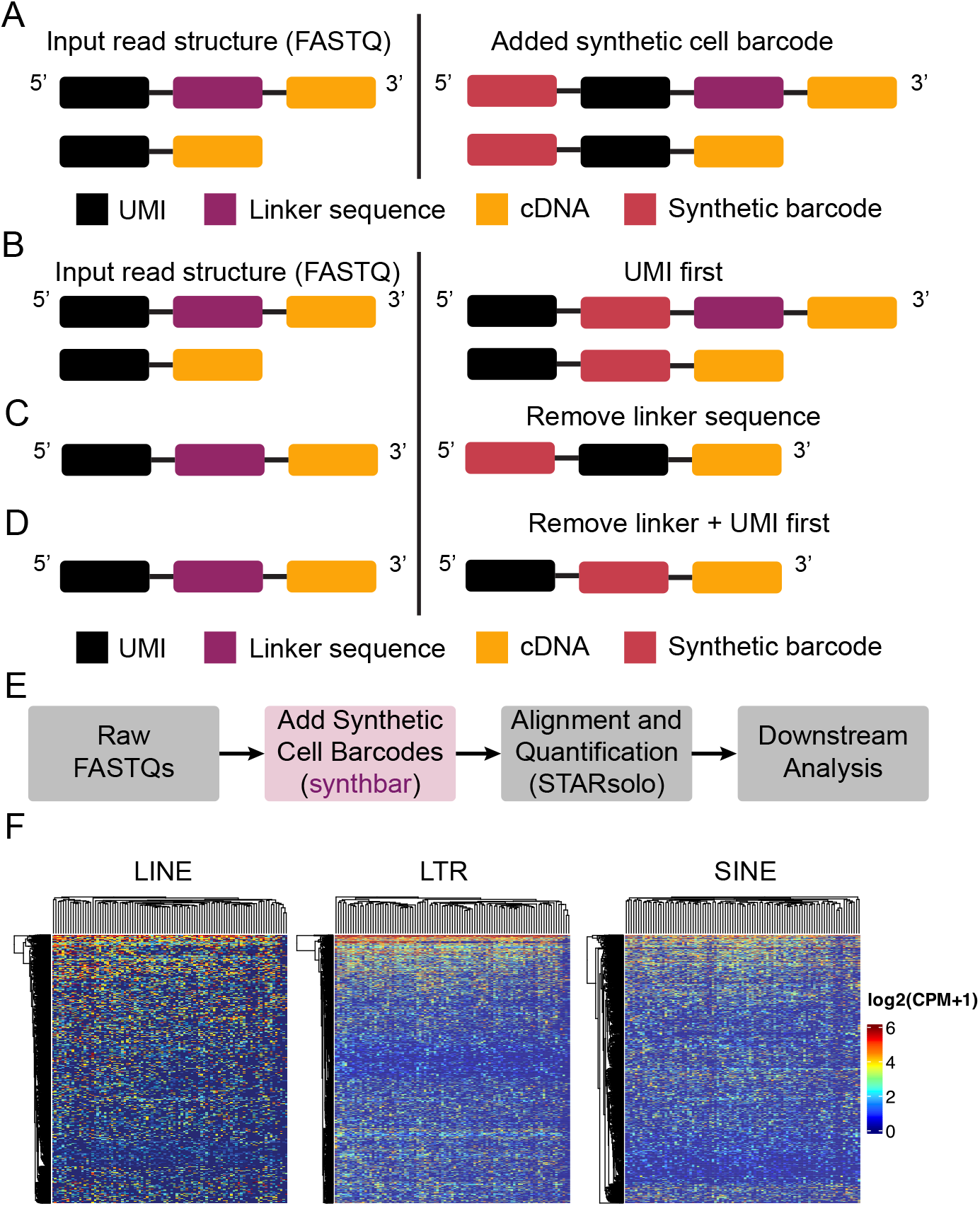
Synthbar is a lightweight tool that can add synthetic cell barcodes and modify standard read structures. **(A)** Default settings for synthbar adds a synthetic cell barcode to the 5’ end of the input read structure across varying read fragment layouts. **(B)** Synthbar can modify the read structure to allow for preservation of the UMI at the 5’ end of the read, while inserting the synthetic cell barcode after the UMI. **(C)** Some library construction methods include a linker or spacer sequence as part of the UMI addition that can be removed, while adding the synthetic barcode to the 5’ end of the read. **(D)** Representative input and output when enforcing the UMI be at the 5’ end of the read, followed by the inserted barcode, removing the linker/spacer sequence. **(E)** Overview of using synthbar in an analysis pipeline. **(F)** Synthbar allows for the probing of transposable elements in K-562 STORM-seq data. TEs shown are intergenic and intronic using Ensembl 101 annotations.

## Example Use Case

In general, synthbar would be run on the raw FASTQ files prior to alignment with tools such as STARsolo or UMI collapsing with UMI-tools (Figure 1E). It can be easily placed into existing analysis pipelines or used to create new analysis pipelines for a variety of scRNA-seq methods, such as Smart-seq-total or Smart-seq3xpress. As a practical use case for synthbar, we have implemented it as part of the analysis workflow for the single-cell total RNA protocol, STORM-seq, which is a paired-end method that incorporates a UMI as part of read 2, but no cell barcode. The lack of a cell barcode makes it inherently unable to work directly with STARsolo and take advantage of the included UMI sequences. As a result, synthbar was used to make a per-cell temporary read 2 FASTQ file that incorporates the default CB at the beginning of the read and meets the expected input read layout for STARsolo. Specifically, this has been implemented as part of the STORMsolo.sh workflow (https://github.com/biobenkj/stormseq_protocols).

Transposable elements (TEs) are repetitive elements in the genome that can act as regulatory elements and are often expressed in a cell type specific manner (1). We applied the STORMsolo pipeline on existing K-562 STORM-seq data to examine long interspersed nuclear element (LINE), long terminal repeat (LTR), and short interspersed nuclear element (SINE) expression in single cells. We reconstruct LINE (n=2540), LTR (n=1588), and SINE (n=1238) expression profiles in K-562 single cells (Figure 1F, Methods).

## Discussion

Cell barcodes (CBs) are essential in droplet-based single-cell analyses for associating sequencing reads, read counts, and other data with individual cells. However, not all scRNA-seq protocols incorporate CBs during library preparation, though they may include UMIs (e.g., STORM-seq, Smart-seq3xpress, or Smart-seq-total). This read structure prohibits the use of tools such as STARsolo that expect a CB as part of the fragment structure in an analysis workflow. Additionally, the presence of a UMI without a CB can increase the number of steps for processing single-cell data and require additional tooling. Although built-in command-line tools can be used to incorporate arbitrary sequences into the read structure, a lightweight and dedicated toolkit designed for this purpose offers advantages by enhancing accessibility, streamlining workflows, and promoting reproducibility. Synthbar was designed and implemented with these criteria in mind. The utility of synthbar extends beyond STORM-seq, including as part of other scRNA-seq protocols like Smart-seq3xpress and Smart-seq-total or single-cell DNA assays such as single-cell whole genome bisulfite sequencing (scWGBS).

Introduction of unique molecular indexes (UMIs) during single-cell library preparation enables the ability to account for PCR duplication bias. It is commonplace for scRNA-seq methods to use UMIs, but less so in other single-cell genomics assays (e.g., single-cell Whole Genome Sequencing [scWGS] and scWGBS). Instead, scWGS and scWGBS methods will de-duplicate based on genomic locations of mapped reads using tools like Picard MarkDuplicates (https://broadinstitute.github.io/picard) or dupsifter (8). Similar to the incorporation of synthetic cell barcodes, it is plausible to imagine a scenario where synthetic UMIs could be added after duplicate marking, in order to use tools that expect a UMI as part of the read architecture. Synthbar is capable of adding synthetic UMI sequences or any arbitrary user-defined sequence. Taken together, synthbar is a lightweight toolkit that supports introducing arbitrary sequences as CBs, UMIs, or other nucleotide motifs in single-cell and bulk sequencing data.

## Acknowledgements

Computation for the work described in this paper was supported by the High Performance Cluster and Cloud Computing (HPC3) Resource at the Van Andel Research Institute.

## Data Availability

Example data to test synthbar is available at https://doi.org/10.5281/zenodo.14646619. K-562 STORM-seq data were downloaded from GSE181544.

## Code Availability

synthbar is available on GitHub under the MIT license: https://github.com/jamorrison/synthbar.

## Methods

### Data download

STORM-seq data were downloaded from GSE296406 and demultiplexed as described in https://github.com/biobenkj/stormseq_protocols.

### Subsampling K-562 STORM-seq

K-562 STORM-seq data were subsampled to 1 million raw reads per cell using seqtk sample (v1.4-r122), setting -s 100. In total, 101 cells were retained for downstream analyses.

### Transposable element expression analysis

synthbar (v0.1.0) was used to add the default synthetic cell barcode (CATATAC) to read 2 of the subsampled K-562 STORM-seq data. The modified reads were then mapped to hg38 (Ensembl 101 and Dec 2013 – RepeatMasker open-4.0.5 – Repeat Library 20140131 intergenic and intronic TE annotations) using STARsolo (v2.7.11a) with modifications: --runThreadN 16, --outSAMtype BAM SortedByCoordinate, -- outSAMattributes NH HI nM AS CR UR CB UB GX GN, --soloStrand Reverse, --soloType CB_UMI_SIMPLE, --soloCBwhitelist None, --soloBarcodeMate 2, --clip5pNbases 0 21, --soloCBstart 1, --soloCBlen 7, --soloUMIstart 8, --soloUMIlen 8, --soloBarcodeReadLength 0, --soloUMIdedup Exact, --soloMultiMappers EM, --soloFeatures GeneFull, and --soloOutFileNames output/ features.tsv barcodes.tsv matrix.mtx. Default parameters were used for all command line options not listed. Transposable elements (TEs) were quantified using https://github.com/huishenlab/TE_quantification_pipeline following the instructions provided in the pipeline’s README. Briefly, Ensembl 101 annotations were used to generate the supporting files (intergenic and intronic TEs) and quantified using the STARsolo BAM files. Cells were then filtered to annotated LINEs, LTRs, and SINEs, keeping loci with at least 20% of cells expressing a given TE in R (v4.4.1). Heatmaps were plotted using ComplexHeatmap (v2.20.0). Post-processing code is available at https://github.com/biobenkj/stormseq_protocols/tree/main/analysis/synthbar.

